# Fluorescent-expressing Modified Vaccinia Ankara Encoding T7 RNA Polymerase

**DOI:** 10.64898/2026.02.22.707307

**Authors:** Nathaniel Jackson, Mahmoud Bayoumi, Luis Martinez-Sobrido

**Affiliations:** Host-pathogen interactions (HPI) and Disease Intervention and Prevention (DIP) programs, Texas Biomedical Research Institute, San Antonio, TX 78227, USA; Virology Department, Faculty of Veterinary Medicine, Cairo University, Giza, Egypt

**Author notes:** Correspondence should be addressed to: Luis Martinez-Sobrido. All experimental work was conducted at USA institutions and/or universities.

**Keywords:** MVA, VSV, RABV, reporter virus, reverse genetics, T7 RNA polymerase

## Abstract

Efficient recovery of negative-stranded RNA viruses from plasmid cDNA requires robust bacteriophage T7 RNA polymerase expression, commonly supplied by Modified Vaccinia Ankara expressing T7 polymerase (MVA T7). Here, we describe the generation and characterization of two fluorescent MVA T7 viruses expressing either enhanced green fluorescent protein (EGFP) or monomeric red fluorescent protein 1 (mRFP1). Both fluorescent-expressing MVA T7 viruses retain plaque morphology, growth kinetics, and transgene expression comparable to parental MVA T7. Importantly, both fluorescent-expressing MVA T7 viruses support rescue of replication-competent recombinant Vesicular Stomatitis Virus (rVSV) while enabling direct visualization of MVA T7 infection. Fluorescent protein expression from MVA T7 viruses facilitates stock generation and titration and allows detection of remaining helper virus during rVSV recovery and amplification. Altogehter, EGFP- and mRFP1-expressing MVA T7 viruses provide a practical tool for monitoring T7-driven rescue of recombinant RNA viruses and identifying helper virus carryover.

**IMPORTANCE:** Reverse genetics systems for negative-stranded RNA viruses rely on robust bacteriophage T7 RNA polymerase expression, commonly provided by a Modified Vaccinia Ankara virus expressing T7 polymerase (MVA T7). However, generation of virus stocks, monitoring MVA T7 helper virus infection and detecting residual virus during rescue of negative-stranded RNA viral can be challenging. We generated fluorescent-expressing MVA T7 variants that enable direct visualization of helper virus infection without compromising viral growth kinetics, transgene expression, or efficient rescue of recombinant Vesicular Stomatitis Virus (rVSV) while enabling direct visualization of MVA T7 infection. MVA T7 viruses expressing fluorescent proteins simplify stock generation and titration, facilitate optimization of negative-stranded RNA rescue conditions, and allow rapid identification of helper virus carryover during generation and amplification of recombinant virus, improving the efficiency, reproducibility, and quality control in reverse genetics systems relying on T7 expression widely used across virology.

## INTRODUCTION

The use of the bacteriophage T7 RNA polymerase has enabled the recovery of negative-stranded RNA viruses entirely from plasmid cDNA. These include the rescue of recombinant Sendai Virus (SeV)(1), Vesicular Stomatitis Virus (VSV)(2), Measles Virus (MeV)(3), Newcastle Disease Virus (NDV)(4), Lymphocytic Choriomeningitis Virus (LCMV)(5), and Rift Valley Fever Virus (RVFV)(6), among others. Expression of T7 RNA polymerase for rescuing recombinant negative-stranded RNA viruses can be achieved using a Vaccinia Virus (VACV) genetically engineered to express the T7 phage polymerase (vTF7-3)(1, 2, 7). However, VACV has several disadvantages, including (i) severe cytopathic effect (CPE) in mammalian cells(8), that may interfere with virus rescue; and (ii) the ability to infect humans(9, 10), necessitating vaccination of laboratory personnel to safely work with the virus. To overcome these limitations, a Modified Vaccinia Ankara (MVA) expressing the phage T7 RNA polymerase was developed (MVA T7)(11, 12). MVA is highly attenuated in mammalian cells due to extensive passaging of VACV Ankara strain in chicken embryo fibroblasts(13), resulting in reduced CPE(8) and improved safety(14, 15). Because of this, MVA T7 has been used to rescue negative-stranded RNA viruses, including, as an example, Rinderpest Virus (RPV)(16), SeV(17), Mumps Virus (MuV)(18), and MeV(19). Efficient viral rescue with MVA T7 requires optimization of viral infection(20), since excessive MVA T7 infection could lead to premature cell death from CPE. Contrarily, insufficient MVA T7 infection could result in inadequate T7 RNA polymerase expression needed for successful viral rescue. Moroever, while MVA T7 infection can successfully be used to generate recombinant negative-stranded RNA viruses, its accumulation during viral rescues can result in MVA T7 carryover contamination.

To address these challenges, we generated two fluorescent–expressing MVA T7 viruses to simplify generation and titration of viral stocks, to enable real-time visualization of MVA T7 infection during viral rescue, and to easily detect the presence of helper MVA T7 during the generation of recombinant negative-stranded RNA viruses. We used homologous recombination of the parental MVA T7 with pRB21 plasmids expressing the enhanced green fluorescent protein (EGFP; pRB21 EGFP) or the monomeric red fluorescent protein 1 (mRFP1; pRB21 mRFP1) to generate MVA EGFP T7 and MVA mRFP1 T7, respectively. These fluorescent-expressing MVA T7 viruses have similar plaque size and morphology in chicken fibroblast (DF-1) cells and in baby hamster (BHK-21) cells to levels comparable to the parental MVA T7 in baby hamster (BHK-21) cells suggesting that insertion and expression of either EGFP or mRFP1 does not affect MVA T7 viral fitness. Importantly, EGFP and mRFP1 facilitate the generation and titration of viral stocks using fluorescent visualization. Importantly, we demonstrate the feasibility of using MVA EGFP T7 and MVA mRFP1 T7 to rescue recombinant Vesicular Stomatitis Virus (rVSV) expressing complementary fluorescent proteins (rVSV mCherry or rVSV EGFP, respectively) with an efficiency similar to that of the parental MVA T7. Notably, expression of EGFP or mRFP1 allowed us to easily determine the presence of residual MVA T7 helper virus during the generation and amplification of rVSV by simply using fluorescent microscopy.

Altogether, our study demonstrates the advantages of using fluorescent-expressing MVA T7 viruses to generate and titrate viral stocks, to monitor levels of viral infection in infected cells, and to ensure the removal of the MVA T7 helper virus during generation and amplification of RNA virus stocks. These fluorescent-expressing MVA T7 viruses enhance the efficiency and reliability of using MVA T7 to generate recombinant negative-stranded RNA viruses.

## RESULTS

### Generation and *in vitro* characterization of fluorescent-expressing MVA T7

To generate fluorescent-expressing MVA T7, we used homologous recombination of the parental MVA T7 with pRB21 plasmids encoding EGFP or mRFP1 under an early/late VACV promoter (pE/L) flanked by complementary sequences downstream of the locus of the VACV VP37 gene (**Figure 1A)**. BHK-21 cells were infected with parental MVA T7 at a multiplicity of infection (MOI) of 1, and after 1 h of viral adsroption, cells were transfected with pRB21 EGFP or pRB21 mRFP1 plasmids. At 24 h post-transfection, robust fluorescence was observed, and cell culture supernatants (CCSs) were collected for subsequent rounds of plaque purification on fresh BHK-21 cells. Individual fluorescent plaques from infected BHK-21 cells were isolated in three sequential rounds to ensure the isolation of a pure population of fluorescent-expressing MVA T7 viruses, lacking any parental MVA T7. Following the final round of plaque purification, EGFP and mRFP1 fluorescent-expressing MVA T7 viruses were amplified in BHK-21 cells to generate viral stocks **(Figure 1B)**.

**Figure 1:**
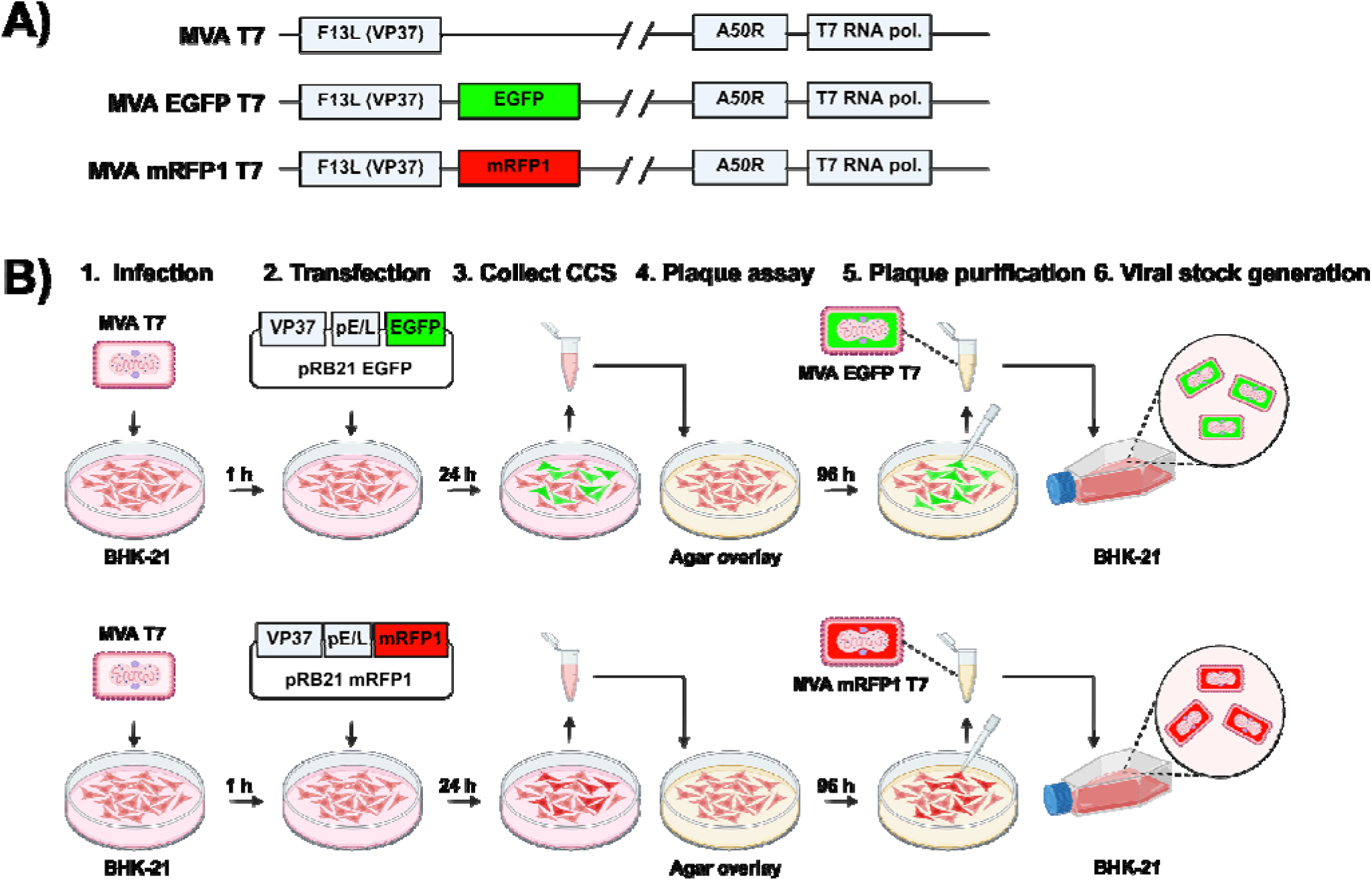
Generation of fluorescent-expressing MVA T7 viruses. **A)** Schematic representation of the genomic organization of parental (top) and fluorescent-expressing EGFP or mRFP1 MVA T7 viruses. **B)** Schematic representation for the generation of fluorescent-expressing MVA T7 viruses by homologous recombination. BHK-21 cells were infected with MVA T7 (MOI = 1) for 1 h, followed by transfection with pRB21 plasmids encoding EGFP (top) or mRFP1 (bottom) under a VACV synthetic early/late promoter (pE/L) and VP37 to facilitate homologous recombination into the genome of MVA T7. After 24 h, cell culture supernatants (CCSs) were collected and used for plaque assays in fresh BHK-21 cells. At 96 h, fluorescent foci were visualized under a fluorescence microscope, and agar plugs were removed, homogenized in cell culture media by vortexing, and used to infect fresh monolayers of BHK-21 cells. This process was repeated three times for plaque purification, and the final passage was used to infect fresh BHK-21 cells in T75 flasks for viral stock generation. Schematics were created with BioRender.com.

To determine whether insertion or expression of the fluorescent proteins into the MVA T7 genome affected viral fitness, we evaluated the plaque phenotype of parental and fluorescent-expressing MVA T7 viruses in DF-1 cells **(Figure 2A)**. Parental and fluorescent-expressing MVA T7 viruses exhibited similar plaque size and plaque morphology, as determine by crystal violet staining **(Figure 2A, bottom)**. Fluorescent-expressing MVA T7 viruses resulted in EGFP- or mRFP1-expressing plaques **(Figure 2A, top)** that corresponded to plaques observed by crystal violet staining. No fluorescence expression was detected in the parental MVA T7 plaques. Importantly, all the viral plaques from MVA EGFP T7 and MVA mRFP1 T7 expressed the fluorescent protein **(Figure 2A)**. To assess whether insertion or expression of the fluorescent proteins impacted MVA T7 replication, multi-step growth kinetics were performed for parental and fluorescent-expressing MVA T7 viruses in BHK-21 cells. Viral titers were comparable among all viruses, peaking between 10 and 10 PFU/mL at 72 h post-infection, with no significant differences observed at 24, 48, or 72 h **(Figure 2B)**. Fluorescence microscopy of the same infected cells at 24 and 48 h post-infection revealed comparable levels of EGFP or mRFP1 expression in cells infected with fluorescent-expressing MVA T7 viruses, whereas no fluorescence was detected in BHK-21 cells infected with the parental MVA T7 **(Figure 2C)**. At 72 h post-infection, all viruses reached 100% cytopathic effect (CPE), resulting in complete cell detachment (data not shown). Altogether, these results confirm EGFP and mRFP1 expression from fluorescent-expressing, but not parental MVA T7, infected cells and that insertion or expreesion of EGFP or mRFP1 does not affect MVA T7 viral fitness.

**Figure 2:**
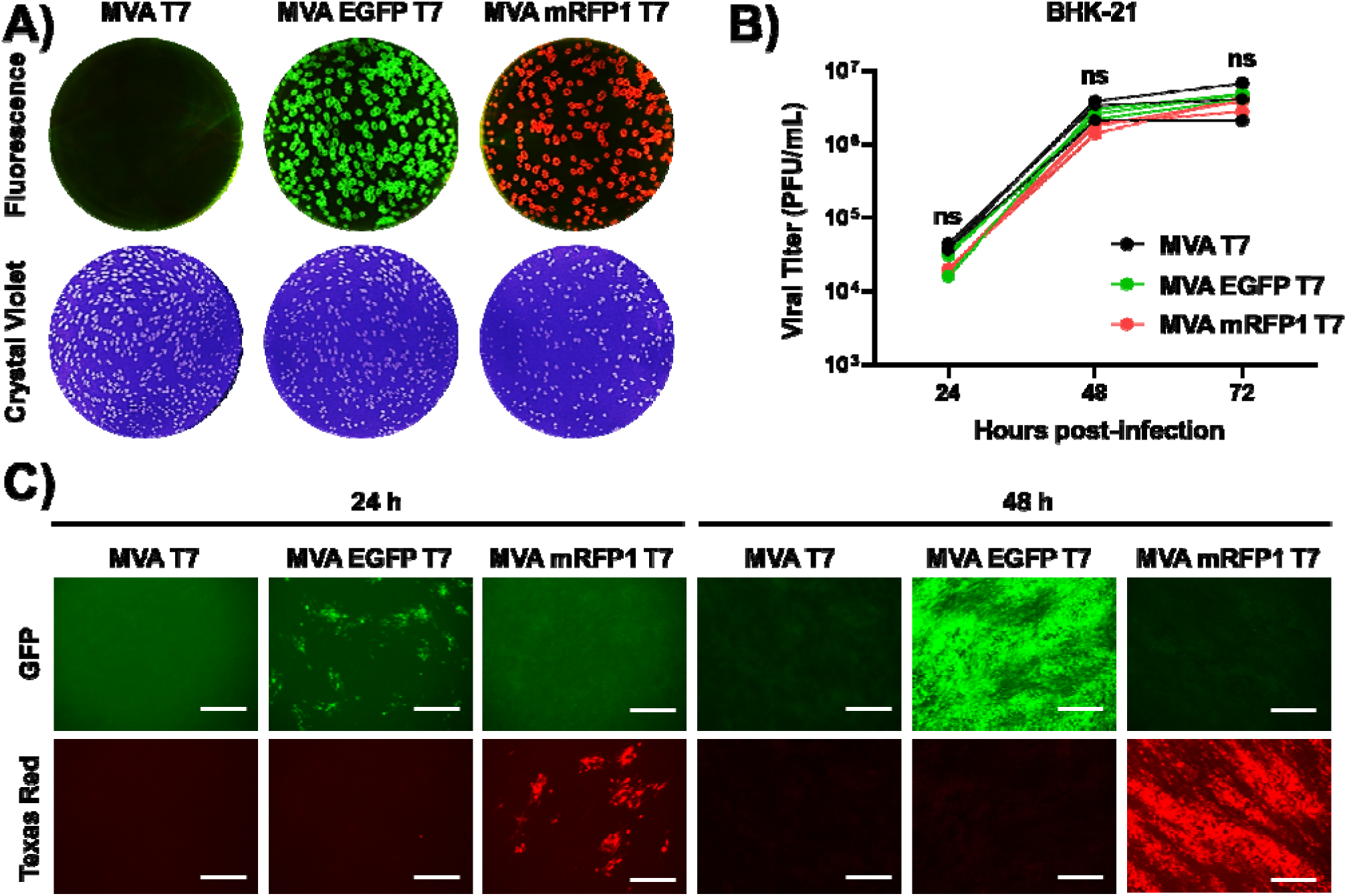
*In vitro* characterization of fluorescent-expressing MVA T7 viruses. **A)** Plaque morphology of parental, EGFP-, and mRFP1-expressing MVA T7 viruses in DF-1 cells after 72 h infection, as shown by fluorescence microscopy (top) and crystal violet staining (bottom). **B)** Replication kinetics of parental, EGFP, and mRFP1 expressing MVA T7 viruses in BHK-21 cells. Statistical analysis was performed using a two-way ANOVA with Tukey’s multiple comparisons test. non-significant (ns) > 0.05 p value. **C)** Fluorescent images of BHK-21 cells infected with MVA T7, MVA EGFP T7, and MVA mRFP1 T7 viruses at 24 h (left) and 48 h (right) post-infection. Scale bar = 300µm.

### Effect of reporter insertion and expression on MVA T7–mediated transient transgene expression

To determine whether insertion or expression of the fluorescent proteins into the VP37 locus of MVA T7 affects transient expression of foreign genes, BHK-21 cells were first infected with parental or fluorescent-expressing EGFP or mRFP1 MVA T7 viruses. Next, infected cells were subsequently transfected with pRB21 reporter plasmids expressing mRFP1 (with MVA T7 and MVA T7 EGFP) or EGFP (with MVA T7 and MVA T7 mRFP1), allowing plasmid-driven fluorescence to be distinguished from MVA T7-encoded fluorescent expression. In all conditions, BHK-21 cells were also transfected with a pRB21 encoding firefly luciferase (Fluc; pRB21 Fluc) to quantitatively measure MVA T7 transient expression. To normalize transfection efficiencies, BHK-21 cells were transfected with a pCAGGS plasmid encoding renilla luciferase (Rluc; pCAGGS Rluc). Transient protein expression was assessed using both qualitative fluorescent and quantitative luciferase readouts (**Figure 3A)**. Fluorescent reporter expression from pRB21 plasmids was comparable in parental and fluorescent-expressing MVA T7 viruses, indicating similar levels of T7 expression (**Figure 3B)**. Consistent with these observations, quantification of luciferase reporter expression from pRB21 revealed no significant differences in Fluc activity in cells infected with parental, EGFP- or mRFP1-expressing MVA T7 viruses (**Figure 3C)**. Together, these results suggest that insertion and expression of fluorescent EGFP or mRFP1 from the VP37 locus of fluorescent-expressing MVA T7 viruses does not affect T7-mediated transient protein expression.

**Figure 3:**
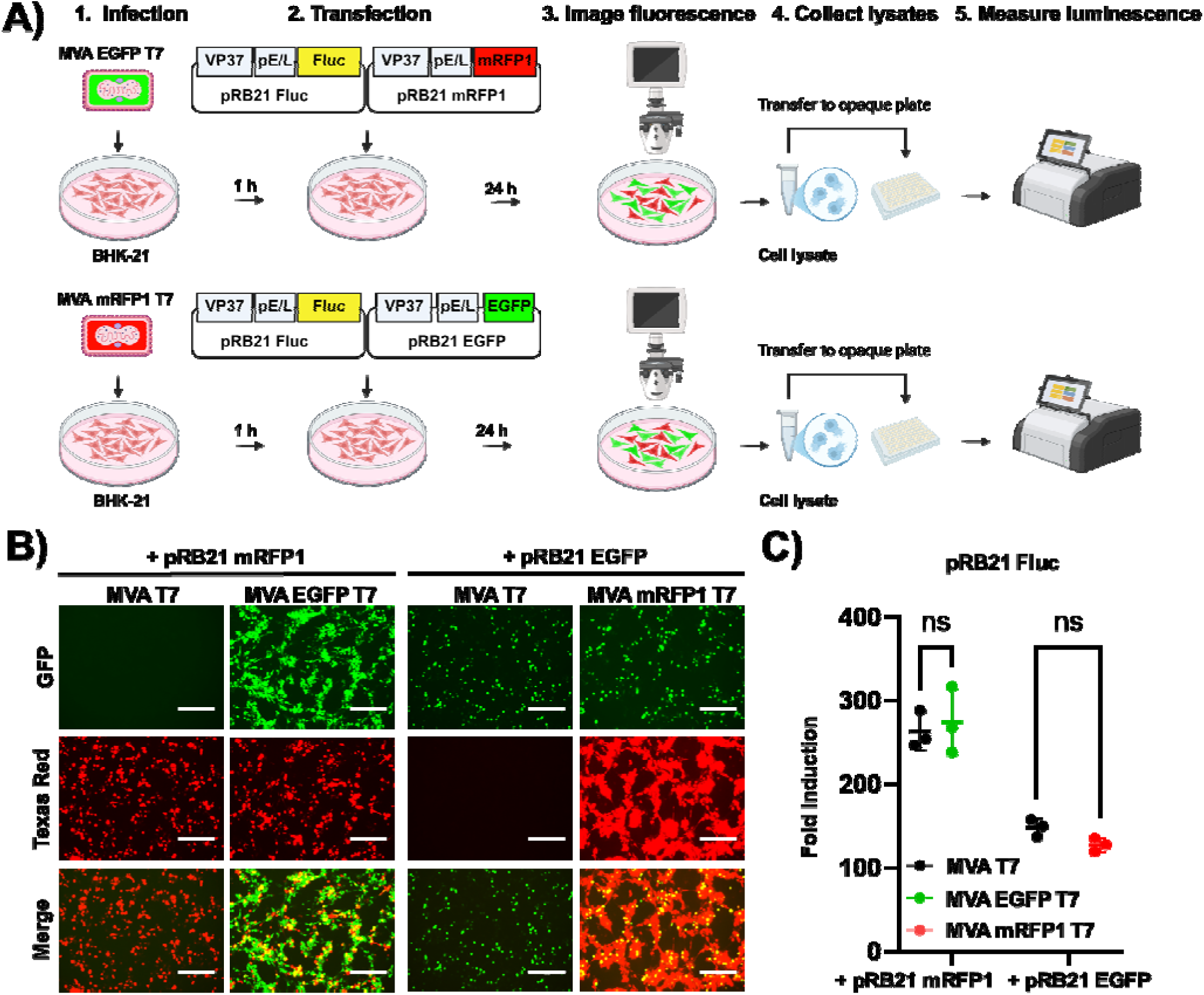
Transient protein expression in cells infected with fluorescent-expressing MVA T7 viruses. **A)** Schematic representation of transient protein expression in cells infected with fluorescent-expressing MVA T7 viruses. BHK-21 cells were infected (MOI=1) with MVA T7 expressing EGFP (top) or mRFP1 (bottom). After 1 h viral adsroption, the viral inoculum was removed and replaced with transfection media containing pRB21 mRFP1 and pRB21 Fluc (top) or pRB21 EGFP and pRB21 Fluc (bottom) plasmids. Cells were also transfected with pCAGGS Rluc to normalize transfection efficiency. Infections with parental MVA T7 were included as control. At 24 h post-infection, cells were imaged by fluorescence microscopy (**B**), and cell lysates were collected to quantify Fluc and Rluc expression using a luciferase plate reader (**C**). Fluc signal was normalized to Rluc expression and fold induction is represented relative to mock-infected transfected cells. Scale bar = 300µm. Statistical analysis was performed using a two-way ANOVA with Tukey’s multiple comparisons test. non-significant (ns) > 0.05 p value. The schematic was created with BioRender.com.

### MVA EGFP T7 driven rescue of replication-competent rVSV mCherry RABV G

To demonstrate the advantages of using fluorescent-expressing MVA T7 viruses to rescue recombinant negative-stranded RNA viruses, we assessed the rescue of a replication-competent T7-driven recombinant vesicular stomatitis virus (rVSV) expressing a fluorescent mCherry and the viral glycoprotein (G) of Rabies Virus (RABV). BHK-21 cells were infected (MOI 1) with parental or MVA EGFP T7 for 1 h, followed by co-transfection of a full-length pVSV encoding mCherry and RABV G together with the corresponding VSV helper plasmids (NP, P, L, and G) under the control of the T7 promoter (**Figure 4A)**. We use the MVA EGFP T7 and the pVSV mCherry RABV G to easily differentiate MVA T7 infected cells (EGFP) from rVSV mCherry RABV infected cells (mCherry). At 24 and 48 h post-transfection, robust mCherry expression from pVSV mCherry RABV G was observed in cells infected with either parental MVA T7 or MVA EGFP T7, indicating successful T7-driven expression of rVSV **(Figure 4B)**. EGFP fluorescence was only detected in BHK-21 cells infected with MVA EGFP T7 and not in parental MVA T7-infected cells **(Figure 4B)**. At 72 h post-transfection, complete CPE was observed for all conditions (data not shown). To demonstrate successful rVSV mCherry RABV G rescue and the lack of MVA T7 helper plasmid in the rescued viral preparation, CCSs were collected and filtered using a 0.2 µm syringe filter and then passaged onto Vero-E6 cells, which do not support productive MVA infection. As a control, collected CCSs were not filtrated and passaged onto Vero-E6 cells. Fluorescence microscopy of infected Vero-E6 cells revealed that, in the absence of filtration (i.e., - Filtration), we could detect EGFP and mCherry, indicating MVA EGFP T7 was retained in the CCSs with rVSV mCherry RABV G **(Figure 4C)**. Contrarily, after two rounds of filtration (i.e., + Filtration), fluorescence microscopy revealed a complete elimination of MVA EGFP T7 while preserving rVSV mCherry RABV G **(Figure 4C)**. As parental MVA T7 does not express a fluorescent protein, residual virus could not be directly visualized by using fluorescent microscopy like with MVA EGFP T7.

**Figure 4:**
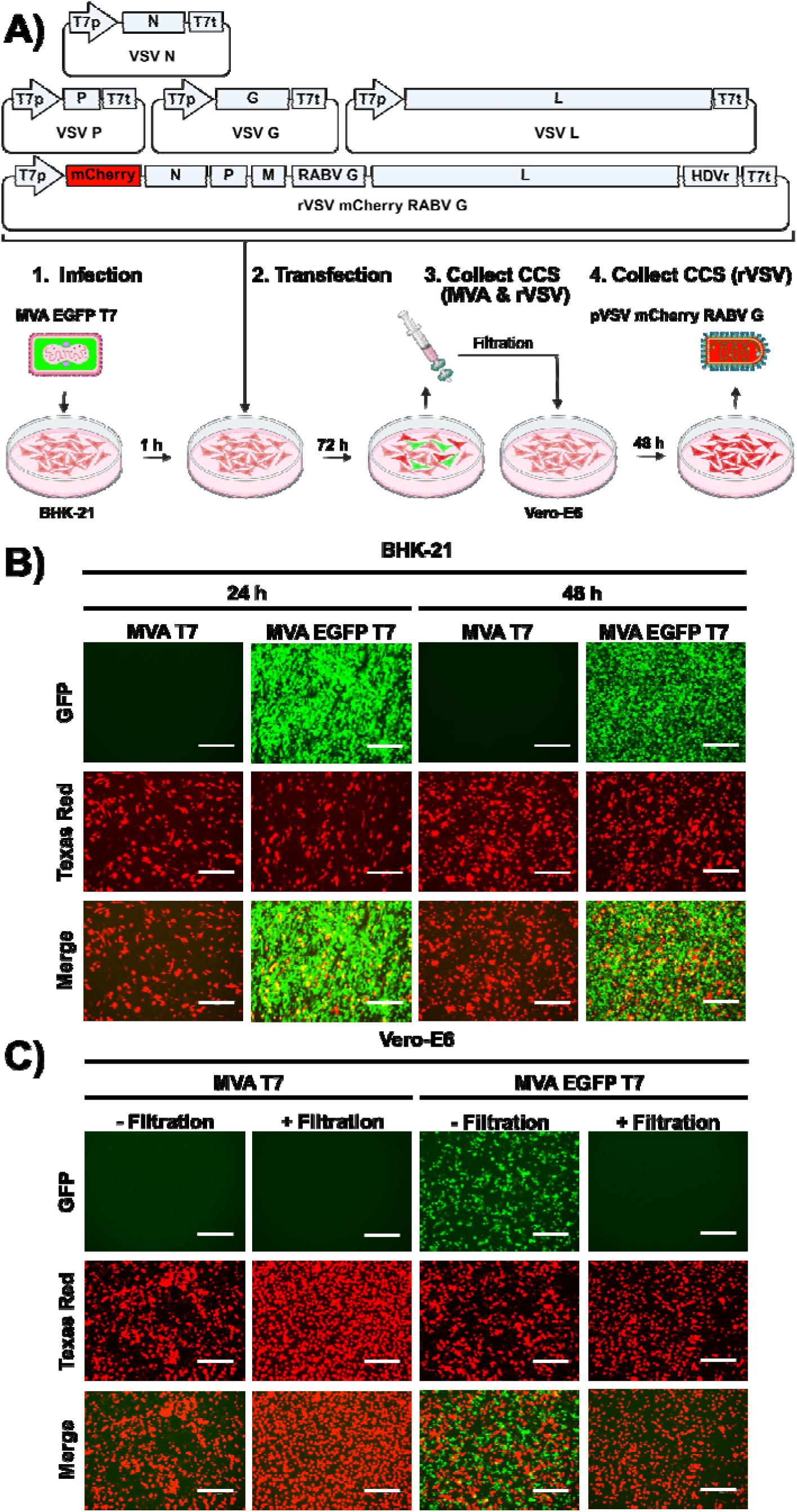
MVA EGFP T7-driven rescue of replication-competent rVSV mCherry RABV G. **A)** Schematic representation of MVA EGFP T7 driven rescue of replication-competent rVSV mCherry RABV G. BHK-21 cells were infected (MOI=1) with MVA EGFP T7. Parental MVA T7 was included as control. After 1 h viral infection, the viral inoculum was removed and replaced with transfection media containing pVSV mCherry RABV G plasmid flanked by T7 promoter/terminator (T7p/T7t) sequences and the Hepatitis Delta Virus ribozyme (HDVr), along with VSV N, VSV P, VSV G, and VSV L helper plasmids under T7 promoter. Fluorescent images were taken at 24 and 48 h post-transfection using a fluorescence microscope (**B**). At 72 h, CCSs were collected and passaged onto Vero-E6 cells and incubated for 48 h. After two rounds of infection, CCSs from Vero-E6 cells were filtered through a 0.2 µm syringe filter. Fluorescent images of unfiltered (- Filtration) and filtered (+ Filtration) CCSs were taken (**C**). Scale bar = 300µm.

### MVA mRFP1 T7 driven rescue of replication-competent rVSV EGFP RABV G

To demonstrate that fluorescent-expressing MVA T7 could support the rescue of T7-driven viruses encoding a more commonly used EGFP, we performed an analogous rescue using an EGFP-expressing pVSV construct **(Figure 5A)**. BHK-21 cells were infected (MOI 1) with MVA mRFP1 T7 and co-transfected with a pVSV EGFP RABV G and the corresponding T7-driven VSV helper plasmids (NP, P, L, and G). Robust EGFP expression was observed at 24 and 48 h post-transfection, indicating efficient transfection with pVSV EGFP RABV G **(Figure 5B)**. These findings correlate with the levels of mRFP1 expression in BHK-21 cells infected with MVA mRFP1 T7 **(Figure 5B)**. Consistent with the results obtained using MVA EGFP T7 (**Figure 4**), complete CPE was observed at 72 h post-transfection (data not shown). Amplification of rVSV EGFP RABV G in Vero-E6 cells combined with filtration, effectively eliminated residual MVA mRFP1 T7 while preserving rVSV EGFP RABV G **(Figure 5C)**. These results demonstrate that fluorescent-expressing MVA T7 viruses efficiently supports T7-driven rVSV rescues while enabling real-time visualization of MVA T7 infection and selective elimination of MVA T7 helper virus as demonstrated with fluorescent microscopy.

**Figure 5:**
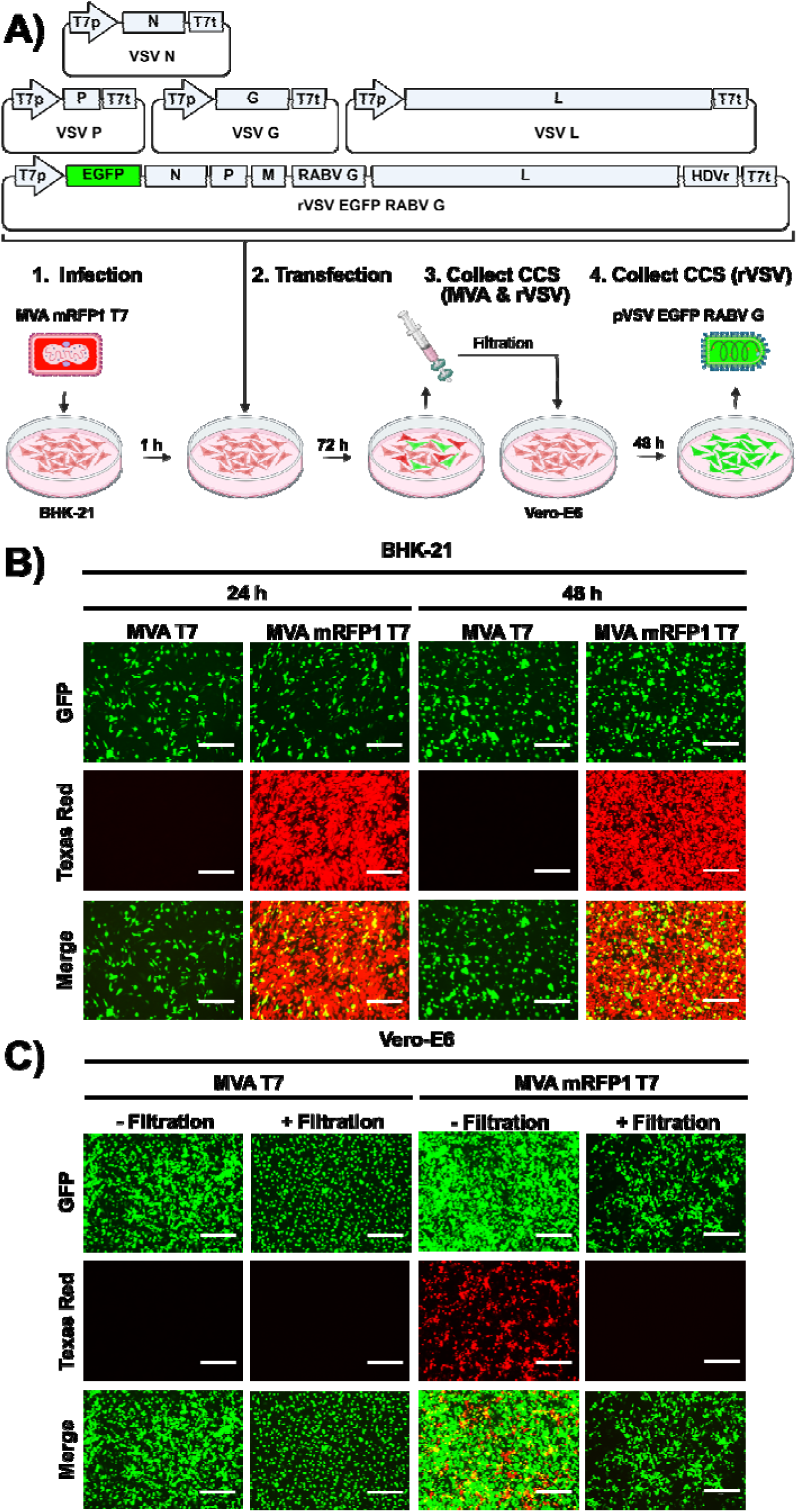
MVA mRFP1 T7-driven rescue of replication-competent rVSV EGFP RABV G. **A)** Schematic representation of MVA mRFP1 T7 driven rescue of replication-competent rVSV EGFP RABV G. BHK-21 cells were infected (MOI=1) with MVA mRFP1 T7. After 1 h viral absorption, the viral inoculum was removed and replaced with transfection media containing the full-length pVSV EGFP RABV G plasmid flanked by T7 promoter/terminator (T7p/T7t) sequences and the Hepatitis Delta Virus ribozyme (HDVr), along with VSV N, VSV P, VSV G, and VSV L helper plasmids under T7 promoter. Fluorescent images were taken at 24 and 48 h post-transfection using a fluorescence microscope (**B**). At 72 h, CCSs were collected and passaged onto Vero-E6 cells and incubated for 48 h. After two rounds of infection, CCSs were filtered through a 0.2 µm syringe filter. Fluorescent images of unfiltered (- Filtration) and filtered (+ Filtration) CCSs were taken (**C**). Scale bar = 300µm.

## DISCUSSION

Rescue of recombinant negative-stranded RNA viruses from plasmid cDNA using MVA T7 has been well documented and established(16–19). However, the use of MVA T7 presents some limitations, including assesing the generation of high titer viral stocks and accurately evaluate MVA T7 viral titers; determing levels of MVA T7 infection during viral rescues; and, evaluating the presence of MVA T7 helper virus from recombinant RNA virus preparations. Here, we describe how we have optimized MVA T7 to incorporate and express fluorescent EGFP or mRFP1, and how we have used these optimized fluorescent-expressing MVA T7 viruses to facilitate the generation and titration of MVA T7 viral stocks, to easily assess viral infection during RNA virus rescue, and, demonstrate the lack of carryover MVA T7 helper virus in recombinant RNA virus preparations. The generation of fluorescent-expressing MVA T7 virus stocks are simplified by monitoring viral-infection time courses using fluorescent microscopy. Likewise, the titration of fluorescent-expressing MVA T7 viruses using plaque assays can be easily achieved by fluorescent microscopy and/or a ChemiDoc system. Importantly, insertion or expression of either EGFP or mRFP1 into MVA T7 does not alter plaque morphology or viral growth kinetics, demonstrating that EGFP or mRFP1 does not significantly impact MVA T7 viral fitness in cultured cells. Extending these findings, we successfully used the fluorescent-expressing MVA T7 viruses to rescue replication-competent rVSV expressing RABV G and fluorescent proteins, which are reciprocal to those present in the MVA T7 helper virus. These results confirm that fluorescent protein choice in both MVA T7 and rVSV does not negatively impact viral rescue efficiency and/or replication. VSV is an exceptionally versatile platform, serving as a vaccine vector(21–25), as a tool to study viral entry(26), and as a safe approach to identify antiviral compounds targeting viral glycoproteins and/or neutralizing antibodies(27–30). In this regard, rVSV expressing reporter genes and the viral glycoprotein of Biosafety Levels (BSL) 3 (e.g. Severe Acute Respiratory Syndrome Coronavirus 2; SARS-CoV-2) or 4 (e.g. Ebola Virus; EBOV) viral pathogens can be safely studied in BSL2 laboratories, greatly expanding accessibility for mechanistic and translational research. More importantly, rVSV expressing the glycoproteins of these BSL3 or BSL4 viruses can be used to safely identify drug-resistant or neutralizing antibody-resistant viruses(27) without the biosafety concerns of conducing these experiments with wild-type SARS-CoV-2 or EBOV.

During optimization of the rVSV rescue using the fluorescent-expressing MVA T7 viruses, we observed that in the absence of filtration of the CCSs, or even following a single round of filtration, MVA T7 could persist despite its attenuation in mammalian cells(8, 14). Potential MVA T7 virus helper contamination in rVSV preparations could represent an important concern for downstream applications, especially those involving the use of VSV as a vaccine. A major advantage of the fluorescent-expressing MVA T7 viruses is the ability to easily monitor the presence of residual MVA T7 helper virus by fluorescence microscopy, enabling rapid confirmation of MVA T7 clearance prior to amplification of the rVSV. This easily monitored feature provides a practical quality-control step that is not readily available with the traditional parental MVA T7. Thus, fluorescent protein expression with these novel MVA T7 viruses mitigates concerns regarding carryover of the helper virus using straightforward visual assessment with fluorescent microscopy. A limitation of our study is that we have only shown the feasibility of using the fluorescent-expressing MVA T7 viruses to rescue rVSV. It remains to be determined the extent of the ability of the fluorescent-expressing MVA T7 viruses to rescue other recombinant negative-stranded RNA viruses.

In summary, we describe the generation and characterization of EGFP and mRFP1 fluorescent-expressing MVA T7 viruses that can be used, similarly to the previously described parental MVA T7, for the rescue of rVSV and potentially other negative-stranded RNA viruses(16–19) that rely on the expression of the bacteriophage T7 RNA polymerase. Importantly, fluorescent-expressing MVA T7 viruses simplify generation and titration of viral stocks and allows easy detection (or lack thereof) of MVA T7 helper virus in rVSV preparation by simply monitoring fluorescent expression using a fluorescent microscope.

## MATERIALS AND METHODS

### Cells

Baby hamster kidney (BHK-21, ATCC CCL-10), African green monkey kidney epithelial (Vero-E6, ATCC CRL-1586), and chicken embryo fibroblast (DF-1, ATCC CRL-12203) were maintained in cell culture medium made up of Dulbecco’s modified eagle medium (DMEM) (Corning) supplemented with 10% fetal bovine serum (FBS) (VWR) and 100 units/mL penicillin streptomycin L-glutamine (Corning), at 37°C in a 5% CO_2_ incubator.

### Viruses and Plasmids

The Modified Vaccinia Ankara virus expressing T7 RNA polymerase (MVA T7) and the pRB21 plasmid were kind gifts from Drs. Bernard Moss at the National Institutes of Health (NIH, Bethesda, MD) and Rafael Blasco at Departamento de Biotecnología, Instituto Nacional de Investigación y Tecnología Agraria y Alimentaria (INIA, Madrid, Spain), respectivelly. Enhanced green fluorescent protein (EGFP) or monomeric red fluorescent protein-1 (mRFP1) were cloned into pRB21 to generate fluorescent-expressing MVA T7 viruses as previously described(31). Similarly, firefly luciferase (Fluc) was cloned into the pRB21 plasmid. A pCAGGS plasmid expressing Renilla luciferase (Rluc) was used to normalize transfection efficiencies. To demonstrate MVA T7-driven rescue of rVSV EGFP RABV G, the pVSV EGFP RABV G plasmid was obtained from Addgene (Plasmid #31833). The pVSV mCherry RABV G plasmid to rescue rVSV mCherry RABV G was generated by cloning mCherry into the XhoI and MscI restriction sites of the pVSV EGFP RABV G plasmid. VSV N, VSV P, VSV L, and VSV G helper plasmids were used for the rescue of the rVSV.

### Generation of fluorescent-expressing MVA T7

To generate MVA T7 expressing either EGFP or mRFP1, confluent monolayers (6-well plate format, 2×10^6^ cells/well) of BHK-21 cells were infected with MVA T7 (MOI = 1). After 1 h, the viral inoculum was removed and replaced with Opti-MEM. Cells were then transfected with 4 µg of pRB21 EGFP or pRB21 mRFP1 using Lipofectamine 3000 (ThermoFisher) according to the manufacturer’s instructions. At 12 h post-transfection, the media were replaced with cell culture media. At 24 h post-transfection, cell culture supernatants (CCSs) were collected for plaque assay, followed by plaque purification. Briefly, confluent monolayers of BHK-21 cells (6-well plate format, 2×10^6^ cells/well) were infected with 10-fold serial dilutions of CCSs collected from transfected cells. After 1 h, the viral inoculum was removed, and an agar overlay was added. At 96 h post-infection, fluorescent foci were observed under an EVOS fluorescent microscope (ThermoFisher). Thereafter, the agar plugs were removed using a P1000 pipette tip, resuspended in cell culture media by vortexing, and clarified by centrifugation for use in subsequent infections. Fluorescent-expressing MVA T7 viruses were amplified in BHK-21 cells and subjected to three rounds of plaque purification. Viral stocks were generated by infecting BHK-21 cells in T75 flasks and collecting and clarifying viral supernatant three days post-infection. Viral stocks were titrated by plaque assay.

### Quantification of parental and fluorescent-expressing MVA T7 viruses

Plaque assays were performed as previously described(32). In brief, confluent monolayers of DF-1 cells (6-well plate format, 2×10^6^ cells/well) were infected with 10-fold serial dilutions of MVA T7, MVA EGFP T7, or MVA mRFP1 T7. After 1 h viral adsorption, the viral inoculum was removed and replaced with cell culture media containing 2% FBS and 0.5% methylcellulose (Sigma-Aldrich). At 72 h post-infection, cells were fixed in 10% formalin for 1 h and imaged for EGFP and mRFP1 expression using a ChemiDoc (Bio-Rad). Next, formalin was removed and cell monolayers were stained with 0.2% crystal violet for 30 min. The plates were washed with water to remove excess stain solution and imaged with a scanner (Epson Perfection V600 Photo Scanner).

### Viral growth kinetics

Confluent monolayers of BHK-21 cells (6-well plate format, 2×10^6^ cells/well, triplicates) were infected (MOI=0.01) with MVA T7, MVA EGFP T7, or MVA mRFP1 T7. After 1 h, the viral inoculum was removed and replaced with cell culture media containing 2% FBS. At 24, 48, and 72 hpi, 150 µL of CCSs were removed and frozen at −80°C. Replicate plates were fixed in 10% formalin at the same times post-infection and imaged under an EVOS fluorescent microscope for EGFP and mRFP1 fluorescent expression using GFP and Texas Red filters. CCSs were titrated in DF-1 cells by plaque assay as described above.

### Fluorescent-expressing MVA T7-driven transient protein expression

BHK-21 cells (12-well plate format, 5.0×10^5^ cells/well, triplicates) were seeded in poly-d-lysine (Sigma-Aldrich)-treated plates 1 day prior to infection (MOI=1) with MVA T7, MVA EGFP T7, or MVA mRFP1 T7. After 1 h, the viral inoculum was removed and cells were transfected, using Lipofectamine 3000 (Thermo Fisher), with 500 ng of pRB21 Fluc together with 500 ng of pRB21 EGFP (for MVA T7 and MVA mRFP T7) or pRB21 mRFP1 (for MVA T7 and MVA EGFP T7), and 250 ng of pCAGGS Rluc to normalize transfection. At 12 h post-transfection, transfection media was replaced with cell culture media and 24 and 48 h after transfection, cells were imaged for fluorescent expression using a EVOS fluorescent microscope. After imaging, cells were collected, lysed with passive lysis buffer (Promega), and assayed for Fluc and Rluc expression according to the manufacturer’s instructions (Promega) using the Synergy H1 plate reader (Agilent). Fluc fold induction was calculated by dividing the Fluc activity by the Rluc activity for each sample and expressed as a result of a fold change relative to the negative control group lacking MVA T7 infection. Data was represented as the average of three biological replicates with SD indicated.

### MVA T7-driven rescue of replication-competent rVSV

Confluent monolayers of BHK-21 (6-well-plate format, 2×10^6^ cells/well, triplicates) were infected (MOI=1.0) with MVA T7, MVA EGFP T7, or MVA mRFP1 T7. After 1 h viral absorption, viral inoculum was removed and replaced with transfection media containing 2 µg of pVSV EGFP RABV G (for MVA T7 and MVA mRFP1 T7 infected cells) or pVSV mCherry RABV G (for MVA T7 and MVA EGFP T7 infected cells) along with VSV N (1µg), VSV P (0.6µg), VSV L (0.2µg), and VSV G (0.4µg) helper plasmids and Lipofectamine 3000 (ThermoFisher). At 12 h post-transfection, transfection media were replaced with cell culture media. At 24 and 48 h post-transfection, cells were imaged for fluorescent expression using an EVOS fluorescent microscope. At 72 h post-transfection, CCSs were collected and centrifuged to remove cell debris. Clarified supernatants were used to infect confluent monolayers of Vero-E6 cells (6-well-plates, 1×10^6^ cells/well) to amplify the rVSV. At 48 h post-infection, CCSs were collected and passaged for a second round of infection in confluent monolayers of Vero-E6 cells. CCSs from amplified rVSV-infected Vero-E6 cells were collected and filtered using a 0.2 µm syringe filter and added to fresh Vero-E6 cell monolayers. This was repeated for an additional round of filtration to remove all the residual parental or fluorescent-expressing MVA T7.

### Statistical analysis

Statistical analyses were determined using two-way ANOVA with Tukey’s multiple comparisons test. Statistical analysis was performed using GraphPad Prism software v.10.6.0 (GraphPad Software, US; www.graphpad.com). The data represent the average of three biological replicates, with the standard deviation (SD). ns: non-significant; p > .05, *p < .01, **p < .001, ***p < .001, ****p < .0001.

## ACKNOWLEDGMENTS

We thank Dr. Bernard Moss for the kind gift of the MVA T7. We also thank Dr. Rafael Blasco for providing the pRB21 plasmid.

